# Defective But Promising: Evaluating Bioinformatic Pipelines for Utility of Defective Interfering RNA Discovery in Plant Viral Infections

**DOI:** 10.1101/2025.05.09.653214

**Authors:** Anthony A. Taylor, Cristina Rosa, Marco Archetti

## Abstract

We explored the utility of the currently available bioinformatics programs ViReMa, DI-tector, DVGfinder, DG-Seq, and VODKA2 for identifying junction points in plant virus high-throughput sequencing (HTS) data that could be tested downstream for antiviral capacity. Specifically, we looked at whether the outputs from these bioinformatic tools generally agree and whether the most frequently identified “defective viral genomes” (DVGs) from these programs are promising defective interfering RNA (DI) candidates for downstream validation. We also explored the possibility of these tools helping us address a larger research question of whether DI RNA are consistently generated and maintained in a specific virus-host combination when conditions are permissive for their replication and accumulation, our “DI prevalence” hypothesis. This was conducted by running eight previously published RNAseq datasets through all five programs and comparing degree of output overlap, most common junction point identified, and whether previously published DI junction points were found. Our results demonstrate a low degree of agreement regarding identified junction points between programs, promise regarding looking at the most commonly occurring junction for DI candidates, and support for our DI prevalence hypothesis. We conclude that bioinformatics workflows have a place in the toolbox of DI and DVG research, but they should not be used alone. We suggest the use of multiple programs on a dataset to better inform decisions regarding deletions to re-create and screen downstream and reiterate the importance of other avenues of evidence in DVG/DI characterization.

## 1. Introduction

### 1.1. What defective interfering RNAs are and how they’ve historically been found

As sequencing becomes more accessible and computational resources more sophisticated, RNA and DNAseq are becoming increasingly attractive options for exploring and analyzing viral sequence diversity. One area of interest to this end involves the profiling of “defective” viral genomes: genomes that are unable to replicate themselves using host cell machinery without complementation in *trans* by other quasispecies members. These defective genomes may play interesting roles in the progression of viral infection and the evolution of viral quasispecies, from stimulation of innate immunity in animals and plants [1,2] to influencing the evolution of multipartism [3]. A subset of these defective genomes that outwardly interfere with the replication of other quasispecies members has aroused much interest, for they are known to lead to amelioration of symptoms, lower viral titer, or differential accumulation of viral proteins [4–6]. Genomes that fall into this category are called defective interfering (DI) genomes, viruses, RNAs, particles, or molecules, and have been known to occur in animal virus quasispecies since the 1950s [4]. In plants, the first record of DIs came from *Alphanucleorhabdovirus tuberosum* (commonly known as potato yellow dwarf virus, or PYDV) in 1983 [7]. Since then, it has been debated if DIs exist in “natural” (non-passaged) infections or if only laboratory passaging generates the required conditions for DI accumulation [8–11]. Laboratory work has concluded that at least for *Tombusvirus lycopersici* (commonly tomato bushy stunt virus, or TBSV), DIs do not require passaging to arise and DIs of a conserved genome organization arise even in infections started with infectious clones [11]. Field work has been published reporting defective RNA found in naturally infected plants [12–16], but the RNA was either not tested for interfering capability or was passaged several times before DIs were isolated and characterized. One field study specifically sought out DIs, but did not find any, concluding that DIs are not found naturally [9], although they leave open the possibility that DIs could exist below the limit of Northern blots’ detection. Determining the natural range of DIs in plant viral infections is important for a robust understanding of viral ecology, for presence of DIs are known to influence disease severity, likely influencing virus-vector-host interactions [15,17,18]. This, coupled with rising interest in DI-based antivirals, warrants the development of effective search methods for locating defective viral sequences in a quasispecies for downstream testing of interfering capability.

The workhorse of DI discovery in plant DI research has historically been Northern blotting, where hybridization with viral UTR-based probes results in blots with bands that do not correspond to known genomic or sub-genomic viral RNA. These bands are then isolated from infected tissue using various methods and tested for dependence on and interference with the full-length virus or its sub-genomic RNAs [19–24]. However, although a reliable old technique, Northern blotting has some disadvantages, including needing to know the sequence of the virus you’re working with to generate the probes and needing a relatively high amount of RNA for detection. RT-qPCR, although simpler and capable of detecting much smaller amounts of RNA, also requires use of primers, meaning that, like Northern blotting, you must know the virus’ sequence and you will only find DIs if those DIs have the genomic region the primers and/or probe bind to.

In response to these drawbacks, a couple recent review papers have suggested that the increase in high-throughput sequencing (HTS) data, along with bioinformatics pipelines to process it, will aid in the identification of novel defective RNAs for downstream analysis [10,25], because many datasets are easily accessible in repositories; only require a computer and coding knowledge to analyze; and contain the entire virome. Furthermore, there are already several bioinformatics tools available that claim to be able to identify defective viral genomes from HTS data: ViReMa [26], DI-tector [27], DG-seq [28], DVGfinder [29], DVG-profiler [30] and VODKA, now replaced with VODKA2 [31].

### 1.2 Brief overview of the bioinformatics pipelines currently available for identification of defective viral genomes

The oldest program in the literature is ViReMa (Viral Recombination Mapper), which was published in 2014 although it is still receiving annual updates. It is built for detecting recombination junction sites, doing so by aligning the 5’ end of a read to a reference genome with a bowtie seed-based alignment specified by the user (default is 25 nucleotides (nt)). It then makes a new segment by either extracting any unaligned nucleotides at the 3’ end of the read or by trimming the first nucleotide from the read. This is done repeatedly until the whole read is either mapped or trimmed. If the whole read is trimmed, it is considered truly unaligned. If the read maps in multiple locations, these locations are noted in output files. The program outputs .txt files with microdeletions, deletions, insertions, substitutions, and recombinations, and has settings allowing the user to specify the length of the bowtie seed used as well as the minimum number of nucleotides that need to be missing to count the read as a recombination junction site. It is designed for short-read data but can be used with long-read datasets if Burrows-Wheeler Alignment (bwa) is specified instead of bowtie in the argument **--aligner bwa** [26].

The second oldest program is DI-tector, which was published in 2018 and uses **bwamem** to align data against host and virus reference genomes before discarding aligned reads and proceeding with unaligned and clipped reads. It then segments each unmapped read into two pieces; “first” and “last,” to map reads that contain junctions. For each read, the first segment’s size will vary from *X* to *N-X* and the last seg from *N-X* to *X* where *N* is the initial length of the read and *X* is the minimum segment length defined by the user with the argument **--Min_Segment** (default = 20). The segmented reads are then aligned to the viral genome of interest using **bwa-aln** and **bwa-samse**. Reads for which the first and/or last segments are unmapped or multi-mapped are discarded, and FLAG values for mapped read segments are processed and compared to cluster the “DVGs” by similar junction location. If two reads map to different reference strands, they are labelled as copy-backs. If two fragments map to the same reference strand, the positions of the two fragments will be compared to characterize deletion/insertion forms of the DVG [27].

Next is DVG-profiler, published in 2019 as part of the High-performance Integrated Virtual Environment (HIVE) platform at the United States Food and Drug Administration. It is not an aligner software and does not accept raw sequencing data, instead accepting pre-aligned and sorted short reads. It counts the number of alignments for each read, and, if it has more than one alignment and the alignments do not have a perfect score based on its scoring algorithm, it inspects the reads in pairs and finds the best-scored pair of alignments that corresponds to a junction. Like DI-tector, it clusters “DVGs” by general location of the deletion, doing so via a peak-detection algorithm, where the breadth of what’s included in the cluster is defined by the user [30].

DG-seq was published in 2020 and differs from the others by being an R script. It imports some functions and then processes user-assigned directories of .bam files. The imported functions consist of a function to import the .bam files and parse them into a data frame for downstream computing, one to create an uncertainty matrix for the reference (which is essentially a map of all short repeat sequences), one to run through the data frame and use CIGAR strings to identify possible junction points, and then a normalization and filtering function that will only keep gaps greater than or equal to a user defined value and are supported by a user-defined number of reads [28].

DVGfinder was published two years later (2022), combining the newest version of ViReMa available at the time with DI-tector. It runs both programs on the input dataset, consolidating the results into a shared output with normalized terminology, noting which reads both programs reached consensus on, and then applying a machine-learning algorithm (trained on *in silico* generated datasets seeded with DVGs) to apply a “probability of being real” value. It does not invent a new search program itself, merely combines two currently available and adds new bells and whistles [29].

Finally, VODKA2 (Viral Open-source DVG Key Algorithm 2) was published in 2023 and replaces the now defunct VODKA [32], which could only detect copy-back DVGs, only screened the last 3,000 base pairs of the reference genome, did not validate via BLAST nor cluster DVG species, and could not run multiple datasets in parallel. It is a set of shell scripts meant to be run in a Docker environment in a specific order. The first step is to create a ‘VODKA2 non-standard viral genome catalogue,’ which is created from the reference genome and is a list of all possible combinations of break and rejoin along with *N* nucleotides upstream of a break and downstream of a rejoin (this process can take up to 48 hours or more depending on the size of the genome). These are written to a large multiFASTA file as they are generated, and a bowtie2 index is then built on this multi-FASTA file and stored for later use. It accepts .fastq input (it’s recommended to trim and gzip them beforehand to help reduce data volume (E. Achouri, pers. commun.)), and then the VODKA2 analysis pipeline can begin, the first step being to use bowtie2 to align the reads against the genome. Then, reads are mapped to the catalogue, junction points are filtered through BLAST to remove all sequences with ambiguous BLAST alignment to the reference genome, junction points are aggregated into a ‘species’ designation, and output tables are generated [31].

### 1.3 Purpose of our meta-analysis

All these programs tout the ability to locate defective viral genomes by applying slightly different methods to identify reads that do not align contiguously to the reference viral genome chosen. However, it is not possible for a computer to distinguish between a read that does not align contiguously because it’s a relevant defective member of the viral quasispecies and a read that has a neutral deletion or is simply a sequencing artifact. Furthermore, sequencing data cannot provide information on interfering capability. As stated previously, HTS datasets have been suggested as boons for DI discovery, largely in the context of DI-based antivirals, by providing more raw material that can be analyzed repeatedly and in a sensitive manner. The thought is to use these programs to identify defective candidates that can be screened downstream for utility as an antiviral: to be effective at this, the programs should have a low false positive rate. Each program has been tested for its tendency to throw false positives and negatives in their respective publications; however, the test datasets are always *in silico* or experimental datasets produced by that lab. We wanted to assess whether, when used on a variety of other publicly available datasets (RNAseq, siRNAseq, infected plant samples from the field, infected plant samples from greenhouses), outputs from different bioinformatic tools generally agree and whether the most frequently identified DVGs from these programs are promising DI candidates for downstream validation.

Furthermore, as stated previously, how common DIs are in non-passaged plant viral quasispecies is still an open question, and HTS datasets, specifically ‘virome’ datasets and the bioinformatic tools used to process them, have been proposed as a resource to identify and characterize (downstream) the defective portion of the quasispecies. Our working hypothesis regarding DI prevalence in virus populations is that, as long as virally encoded proteins can assist genome replication in *trans* and that these *trans-*assisting virally encoded proteins increase the fitness of the genome they help replicate in a non-linear manner, DIs should be present in the viral quasispecies. This leads to the prediction that if DIs are found in one instance of a virus/host combination, then the conditions for DI generation and accumulation --- whatever they are --- are permissive and we should still be able to locate DIs of that specific genome organization in any other instance of the same combination of virus species/host species. We tested this hypothesis by choosing three datasets where the host/virus combination has literature-backed canon DIs and three where the virus is known to produce DIs in other hosts but has not been assessed for DIs in the sequenced host.

To explore these two questions --- whether (and which) bioinformatic tools show promise for downstream DI characterization and whether our working hypothesis on DI prevalence has any virome data support --- we performed a meta-analysis utilizing eight previously published datasets: three of a host/virus combination known to produce DIs, three of a host/virus combination where the virus is known to produce DIs in other hosts but the DI status of the sequenced host is unknown, and two controls, RNAseq of SARS-CoV2-infected cells treated with a DI antiviral and one *in silico* generated RNAseq dataset seeded with random defective viral genome reads. Our results demonstrate a low degree of agreement regarding identified junction points between programs, including little agreement on the most commonly occurring junction point. However, the junction point(s) identified as ‘most common’ were typically large, disruptive deletions, meaning that they could be explored downstream for antiviral capability with the caveat that this is just a single line of *in-silico* evidence. Furthermore, we did find support --- although not strong - -- for our DI prevalence hypothesis, with two out of three “known combination” datasets sporting canon junction sites from other papers.

## 2. Materials and Methods

### 2.1 Selection of datasets

The NCBI SRA database was searched for datasets of plants infected by each virus known to generate DVGs, resulting in 12 possible viruses to choose from: *Tombusvirus cymbidii* (Cymbidium ringspot virus, CymRSV), *Betacarmovirus brassicae* (turnip crinkle virus, TCV), *Pomovirus solani* (potato mop-top virus, PMTV), *Nepovirus nigranuli* (tomato black ring virus, TBRV), *Potexvirus flavitrifolii* (clover yellow mosaic virus, CYMV), *Curtovirus betae* (beet curly-top virus, BCTV), *Begomovirus manihotis* (African cassava mosaic virus, ACMV), *Begomovirus manihotisafricaense* (East African cassava mosaic virus, EACMV), *Cucumovirus CMV* (cucumber mosaic virus, CMV), *Orthotospovirus tomatomaculae* (tomato spotted wilt virus, TSWV), *Tobravirus tabaci* (tobacco rattle virus, TRV), and *Bromovirus BMV* (brome mosaic virus, BMV)). Three datasets with literature-backed host-virus combinations were picked: CMV in *Nicotiana tabacum*, BMV in *Hordeum vulgare*, and CymRSV in *Nicotiana benthamiana* (Table 1). Then, three test datasets containing viruses that are known to produce DIs on other hosts, but which the sequenced host has not been tested, were selected. These include TCV in *Cicer arietinum*, CYMV in *Verbena officinalis*, and TSWV in *Solanum lycopersicum* (Table 1). For positive controls, a synthetic dataset seeded with DVGs of known sequence was used [29], as well as two datasets (unpublished) consisting of SARS-CoV2 infected human cells treated with DVGs of known sequence produced by in vitro transcription. Note that the viral references used for CYMV and TSWV are not the references used in the publication, since for CYMV there is no publication associated with the SRA data and for TSWV the exact isolate of TWSV used in the paper has not been sequenced. An isolate of TSWV that has previously been found to infect *S. lycopersicum* as a reference was therefore used.

**Table 1.** Information regarding datasets used in this meta-analysis. ‘Breakpoint’ refers to the nucleotide on the viral reference genome where the internal deletion begins, and ‘re-initiation point’ refers to the nucleotide on the viral reference genome where the internal deletion ends.

| Virus | Host | SRA Run Accession Number(s) [Publication] | Viral reference (GenBank) | Paper(s) supporting DIs in this virus + host | Papers(s) supporting that this virus generates DIs | Junctions used in Analysis (breakpoint/re-initiation point) |
| --- | --- | --- | --- | --- | --- | --- |
| *Cucumovirus CMV* (CMV) RNA3 | *Nicotiana tabacum* | SRR6374495 | AB008777 | [33] | [33] | 311/621 |
|  |  | SRR6374496 [30] |  |  |  | 330/487 |
| *Bromovirus BMV* (BMV) RNA3 | *Hordeum vulgare* | SRR10119525 [34] | NC_002028.2 | [35,36] | [35,36] | 369/868 |
|  |  |  |  |  |  | 201/266 |
|  |  |  |  |  |  | 366/865 |
| *Tombusvirus cymbidii* (CymRSV) | *Nicotiana benthamiana* | SRR039711 [37] | NC_003532 | [2] | [2] | 164/1351 |
|  |  |  |  |  |  | 1463/4327 |
|  |  |  |  |  |  | 164/1347 |
|  |  |  |  |  |  | 1504/4347 |
|  |  |  |  |  |  | 4407/4612 |
| *Betacarmovirus brassicae* (TCV) | *Cicer arietinum* | SRR5571020<br>SRR5568966 [38] | NC_003821.3 | N/A | [21] | 141/3863 |
| *Potexvirus*<br>*flavitrifolii*<br>*(CYMV)* | *Verbena*<br>*officinalis* | SRR11928841<br>SRR10490212<br>[No pub.] | NC_001753.1 | N/A | [22] | 757/6603<br>757/6681<br>814/6788<br>831/820 |
| *Ortho-*<br>*tospovirus to-*<br>*matomaculae*<br>*(TSWV)*<br>*Segment L* | *Solanum*<br>*lycopersi-*<br>*cum* | SRR12953073<br>[39] | OM902668.1 | N/A | [20,23,24,40,41] | 1031/6506<br>74/5681<br>138/5930 |
| SARS-CoV2 | *Homo sa-*<br>*piens* | Dataset (experi-<br>mental) provided<br>by M. Archetti<sup>3</sup>,<br>unpublished | NC_045512 | DVGs are<br>synthetic | N/A | 781/19664<br>20340/29902 |
| *Potyvirus ra-*<br>*pae (TuMV)* | *Brassica*<br>*rapa*<br>*subsp.*<br>*rapa* | Dataset (*in silico*<br>generated) in sup-<br>plementary mate-<br>rials of [29] | GitHub of [29] | DVGs are<br>synthetic | N/A | Long list, provided in the supplemen-<br>tary materials of [29] |

### 2.2 Workflow and software

All computing work was done on the ROAR Collab (RC) supercomputer at Penn State University on the ‘open’ account with standard cores, 1 node (4 cores per node), 1 core per task, and 200Gb memory in May-June 2024. Bioinformatics software were obtained from the following sources: https://www.sourceforge.net/projects/virema (ViReMa v0.29), https://www.di-tector.cyame.eu (DI-tector v0.6), https://rnajournal.cshlp.org/content/26/12/1905/suppl/DC1 (DG-seq, run on RStudio 2024.12.1+563 “Kousa Dogwood” running R version 4.3.2 (2023-10-31 ucrt) -- “Eye Holes”), https://github.com/MJmaolu/DVGfinder (DVGfinder v2), and https://github.com/lopezlab-washu/VODKA2 (VODKA2 (2.0)). DVG-profiler [30] requires use of the HIVE software as described in its publication, which at the time of writing appears to be retired. The source code (https://github.com/kkaragiannis/DVG-profiler/) could not be run on RC, so DVG-profiler was dropped from the study.

Across all datasets, ViReMa was run with a seed of 20nt (--Seed 20), except for TCV and CymRSV, which had a seed of 5nt due to those datasets having reads of 35nt or less, and TSWV, which had a seed of 25nt due to reads being 300nt long. Other ViReMa settings were the use of 4 processors for parallel processing (--p 4) and only counting deletions with a deletion length of 5 or greater (--MicroInDel_Length 5). All other settings were default. DI-tector was set to show only junctions with count reads greater than 5 (-n 5), to skip alignments with indels smaller or equal to 5 (-l 5), and to run on 4 threads for parallel processing (-x 4), with all other settings default. DVGfinder was also set to run on 4 threads (-n 4), with all other settings default. VODKA2 was run as the scripts on GitHub (https://github.com/skybird99-anthony-taylor/DIs_in_RNAseq_data) indicate. DG-Seq required some tweaks to run on R v4.3.2, and the edited file with all its annotations is available on GitHub with the other scripts. All programs were run in a Miniconda3 environment ‘bioinformatics_env’ which had all programs and program dependencies (as indicated by ReadMe files or error messages) downloaded into it via **conda install**.

As a control, bwa-mem version 0.7.18-r1243-dirty was installed to the ‘bioinformatics_env’ environment using **pip**. Bwa was selected due to the use of a dataset with long reads (the SARS-CoV-2 dataset), since the upper limit for read length for bowtie is ∼1,000bp, as described in the bowtie user manual. Bowtie2 does not have an upper limit, however, the only program to use bowtie2 was VODKA2, and so bwa was selected as the more relevant program to use. Datasets were downloaded off the SRA database using NCBI’s sra-toolkit (v3.1.0, centos linux 64 bit) into a ‘datasets’ sub-directory, and viral reference genomes corresponding to the SRA data’s publications (see Table 1) were also downloaded. All SRA datasets were run through the **fastqc** command on default settings to check for quality, with all run quality scores >32 excepting reads from the SARS-CoV2 datasets, which were generally between 2 and 30. If datasets were not pre-trimmed, adaptor sequences and primers were trimmed using **cutadapt** (default settings) based on the ‘overrepresented sequences’ tab in the fastqc report. SRA datasets were then aligned to the RefSeq genome and transcriptome of the plant host using **bwa-mem**, with the fully processed output zipped as a .fastq.gz file for faster downstream processing. Control datasets consisted of *in vitro* transcribed RNA of a SARS-CoV2 DI [25] transfected in human cells (one dataset from cells before passaging and the other after three passages) as well as an *in-silico* generated dataset with 72 random DVGs of *Potyvirus rapae* (turnip mosaic virus, TuMV), each present in a proportion of 0.003 in regards to the total number of reads (which included both TuMV and reads from the ‘host,’ *Brassica rapa* subsp. *rapa*).

### 2.3 Analysis

For known virus/host combinations, junctions used for analysis were taken from the supporting paper (Table 1). For unknown virus/host combinations, junctions used for analysis were taken from papers with sequences reported for that virus’ DIs. This was citation [21] for TCV, [22] for CYMV, and [23] for TSWV. Considering that the junctions that make up a particular virus’ DIs are often heterogeneous, a breakpoint and/or re-initiation point shift of up to 5 nucleotides in either direction was tolerated to count a junction as a ‘positive’ result. R scripts were written to search through the large output files for junctions, with the 5nt wiggle room. Considering that all published DI sequences used were deletion DIs, in programs that gave multiple DVG outputs (e.g., insertion, deletion, copy-back) only deletions were analyzed. R code for analysis can be found at the project’s GitHub. DGseq is an R program already, so R code was not written for this analysis: however, as previously stated it required some bug fixing to run, and that code can be found on the GitHub. VODKA2 was originally created to run in a Docker environment and does not function properly without Docker: also provided in the GitHub is an example .txt file of code with instructions detailing how to edit and run the VODKA2 scripts to get it to function in a non-Docker environment. These fixes were a mix of those discovered during analysis by A. Taylor and those suggested by VODKA2’s author, E. Achouri. For the manual alignment using **bwa-mem**, .fasta reference files were generated containing the sequence across the junction reported in their respective papers for all junctions investigated. Samples in .fastq.gz format were aligned against the junction sequence using default settings, filtered to remove all unmapped reads (-F 4), and had the results analyzed. Note that this method of analysis makes it unnecessary to indicate ‘total junctions’ and ‘unique junctions’ as done for the other programs because reads are being directly aligned to a junction point and so will only output reads that have that junction. Junction reference files, except for the COVID-19 DI, are available on the previously linked GitHub.

For determination of junction overlap across the programs, compiled datasets were created taking all the BP and RI values provided in the programs’ outputs and compiling them in a new .csv file with a third column of ‘program’ indicating which program’s output that junction was from. A fourth column of ‘ID’ was created whereby the BP and RI from each row was combined into one character separated by an underscore (e.x. BP of 300 and RI of 3,381 would become ID 300_3381). These combined datasets were then imported to R and visualized as UpSet plots using the package ComplexUpset.

For investigation into the most frequently reported junction in the program outputs, R codes were written to count all instances of a particular junction in the output and then list them in descending order. This list was then manually evaluated and binned to create final instances counts (because using this method a junction point of 1456/3400 and 1457/3408 would be considered separate, even though they’re within the 10nt wiggle room). This instance count was then divided by the total number of DVGs found by the program for that dataset to provide a percent-of-output value. The deletions caused by the most reported junction points were then re-created on SnapGene version 7.2.1 and the effect on ORFs and protein amino acid sequences were noted.

## 3. Results

### 3.1. Canon DI junction sites were not consistently found by bioinformatics programs

The first question we explored was whether canon junction sites reported in previous papers could be found in other RNAseq datasets of the same host/virus combination. Table 2 summarizes this search in the eight analyzed datasets. Of the three virus/host combinations that have been previously shown to produce DIs (“known” combinations), only two had previously published DI junctions identified in the RNAseq datasets used: BMV in *H. vulgare* and CymRSV in *N. benthamiana*. BMV had one junction (201/266) identified by DI-tector, ViReMa, DVGfinder, DG-seq, and bwa-mem, while CymRSV junctions were only found in ViReMa. Only one of the RNAseq datasets involving a known DI-producing virus in a novel host (“unknown” combinations) had a junction corresponding to previously published DIs from that virus, and this was done by ViReMa in TCV, where the program identified antisense deletions corresponding to the known junction site 141/3863. A second junction of BWV, 875/362, was reported by several programs but was outside of the 10nt wiggle room and so was not counted as a valid found junction as per our methods.

Considering the control datasets, the TuMV dataset that was *in silico* generated saw all programs find junctions, however, all programs except BWA-MEM found far fewer than the 72 junctions purposely seeded into the dataset. VODKA2 found the fewest seeded deletions, at only 8. DVGfinder and DG-seq found more junction sites than were seeded, meaning that 10 and 5 of those junction sites were false positives, respectively. TuMV results are not listed in table 2 due to length but can be found in supplemental table 1. Bwa-mem found all 24 with no false positives. The SARS-CoV-2 dataset saw no programs except BWA-MEM find the two junctions present in the synthetic DIs.

**Table 2.**
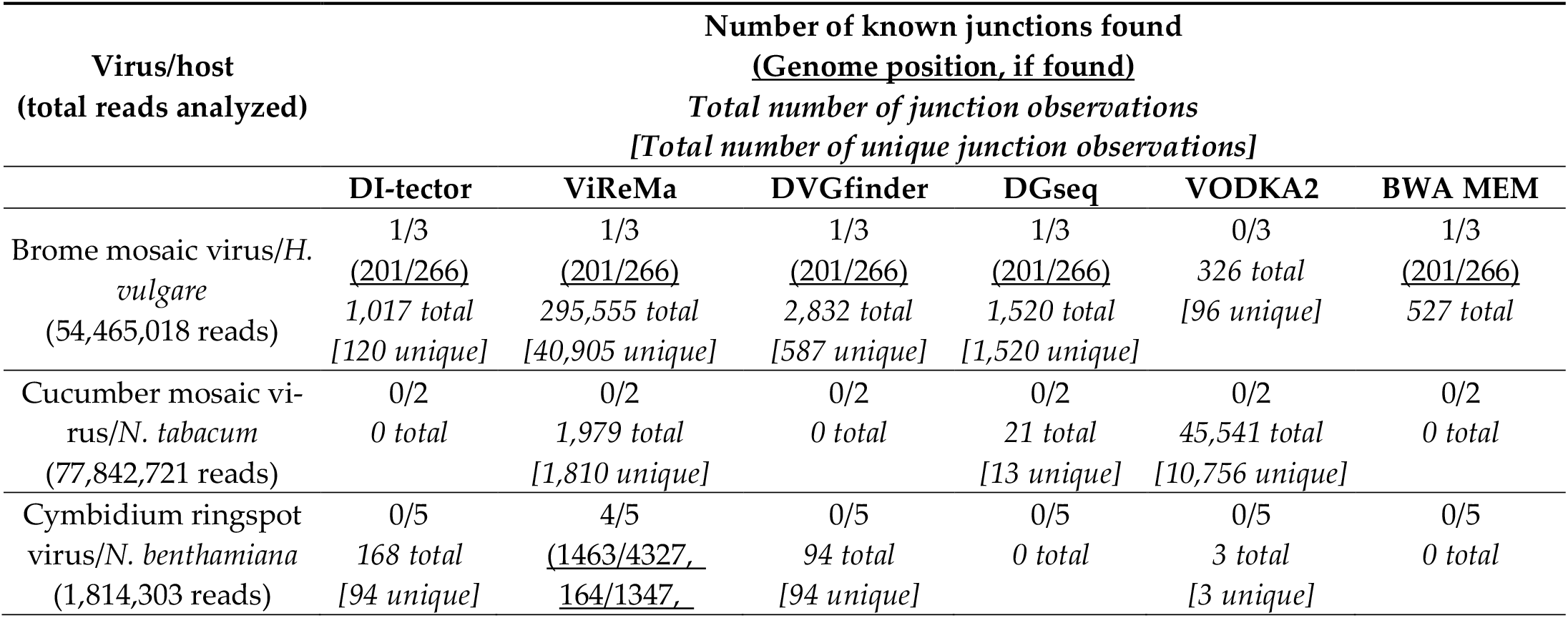
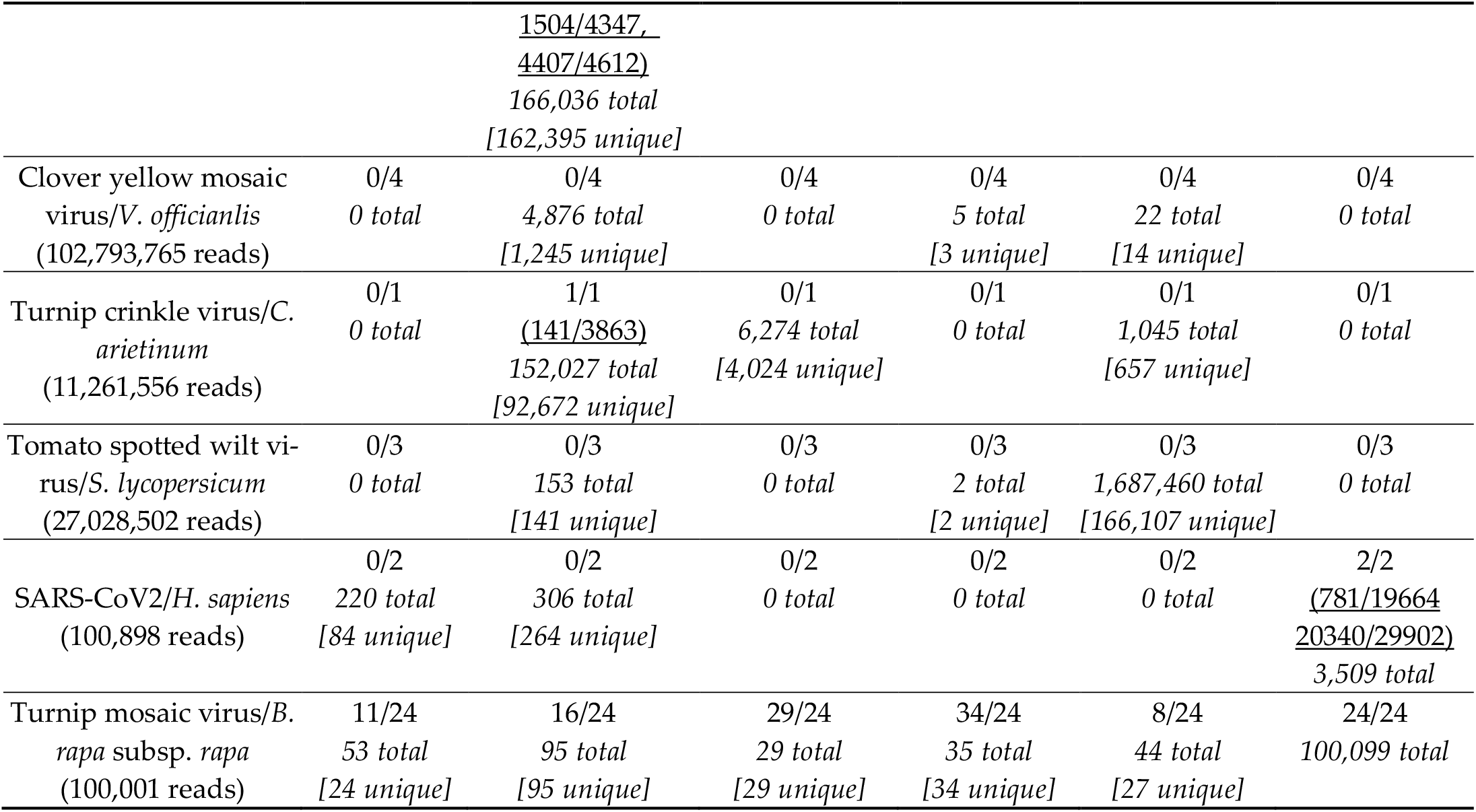
Table detailing the number of canonical junctions found for each virus/DI pair from each program (e.g., 1/3), which canonical junction (if any) were found (e.g., 201/266), total number of output junctions (canonical or non-canonical) provided by the program (e.g., *1,017 total*), and total number of unique junctions (e.g., *[120 unique]*).

In sum, **table 2** demonstrates that two out of three known DI-producing host/virus combination datasets saw at least one bioinformatics program find canon DIs, while only one out of three of the unknown DI-producing host/virus combination datasets, TCV in *Cicer arientum*, had a canon junction point identified by ViReMa.

### 3.2 There is little overlap in results between programs

**Figure 1** demonstrates the overlap in output between all the programs for each dataset. ViReMa and VODKA2 have the highest amount of output, numbering in the tens to hundreds of thousands of unique junction points. Despite this, there is always very little overlap between the two programs. There are two other main points of interest: one, that in four of the six datasets DI-tector identified no deletions and so is not present (Fig. 1D, E, F, and G); and two, that in two cases, BMV and TCV (Fig. 1C and F), DVGfinder identified junctions that were not found in either DI-tector or ViReMa, which is unusual considering that DVGfinder’s search algorithm is made up of those two programs. In the case of TCV, DVGfinder ascribes these junctions to ViReMa, for it records no instances of a DI-tector identification for any of its output junctions. In the case of BMV, DVGfinder ascribes some of these junctions to ViReMa, some to DI-tector, and some to both. Therefore, our results show that in some cases DVGfinder writes junctions to output that are not found by ViReMa or DI-tector when those programs are run independently. **Figure 1** also demonstrates that in most datasets there were no junctions found by all programs, except TCV and the control TuMV. For TCV, all programs that gave results for this dataset find 692/1234, which fuses ORFs 1 and 2, and 1616/2191, which truncates ORF2. For TuMV, consensus junctions were 6000/6040, 3859/7242, 1784/6060, 3516/5147, 1909/8273, and 1051/2302: all true positives. None of these junctions were the most common junction found in the datasets as described in **table 3**, except for 6000/6040 of the TuMV dataset.

**Table 3.**
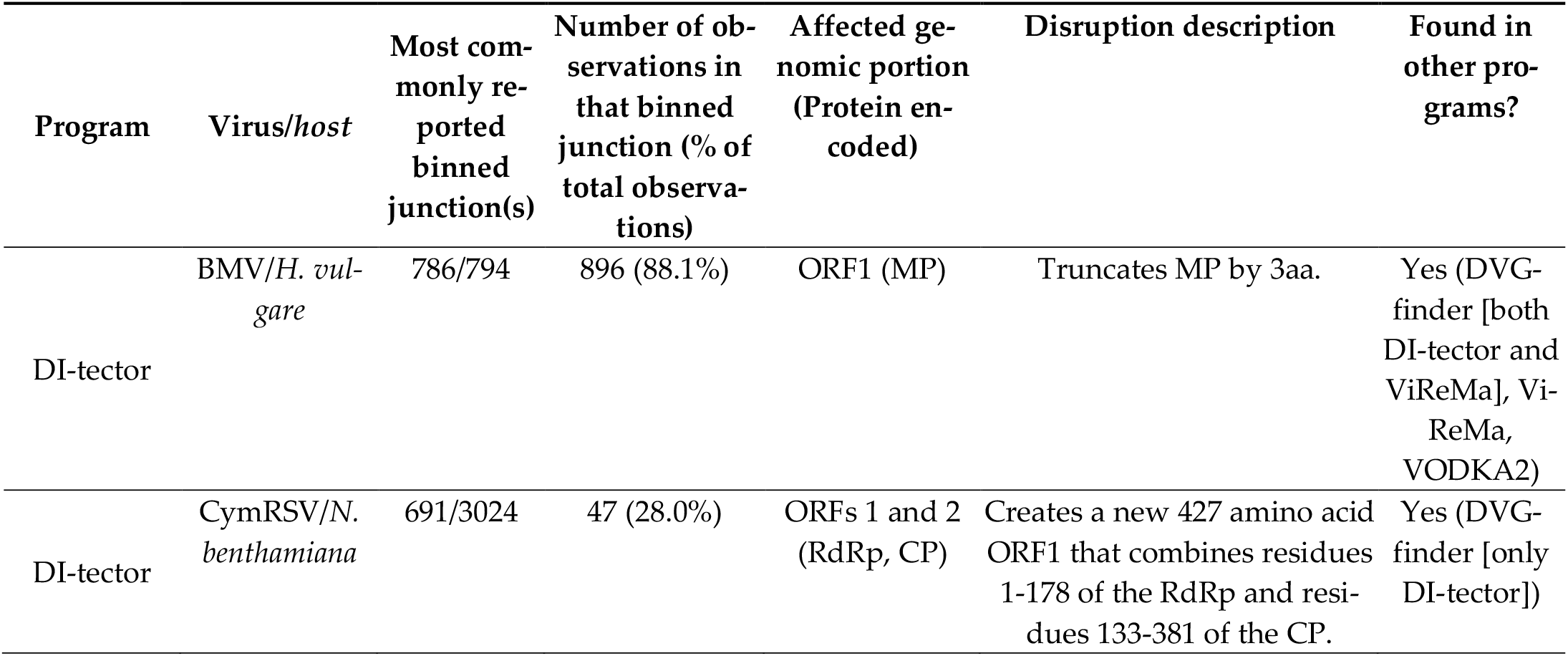
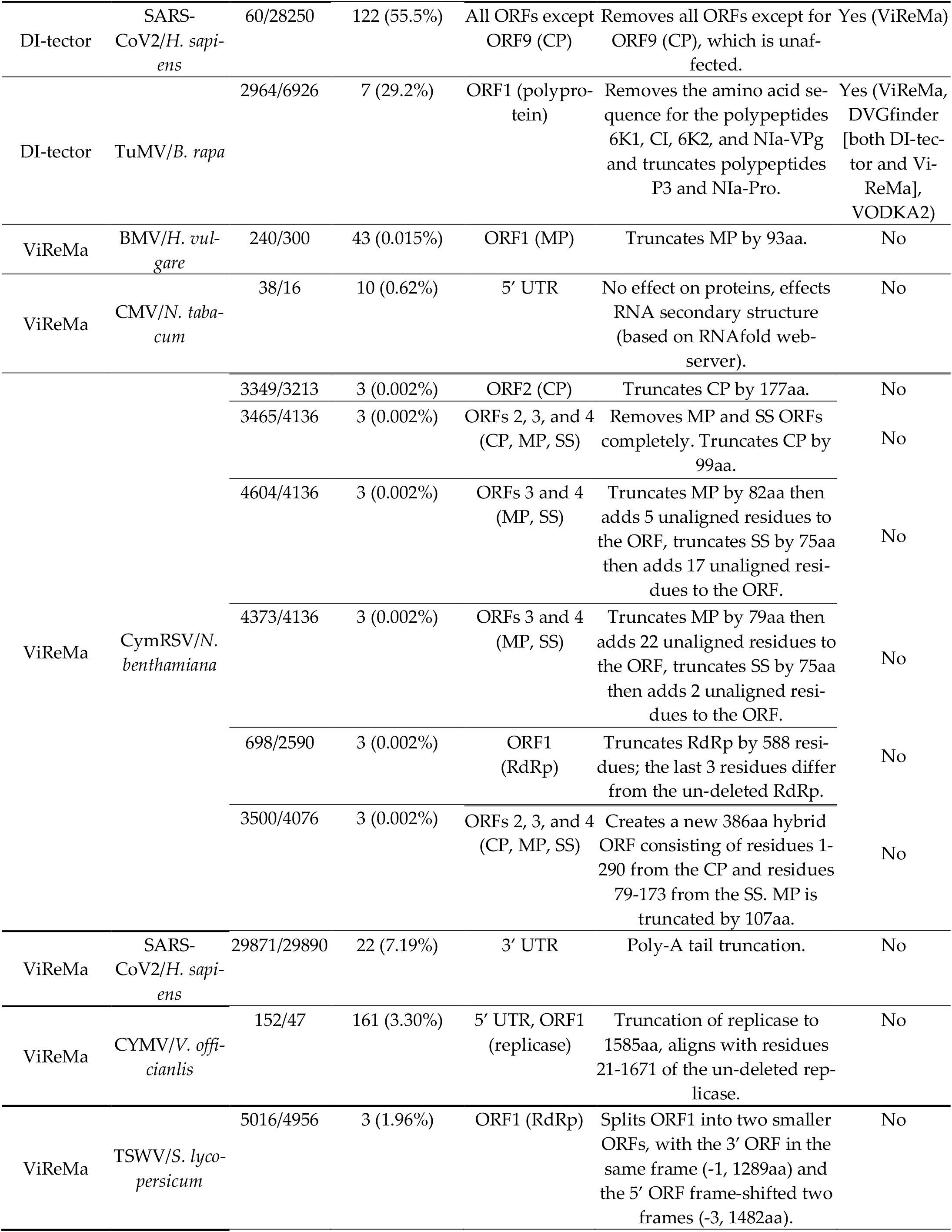
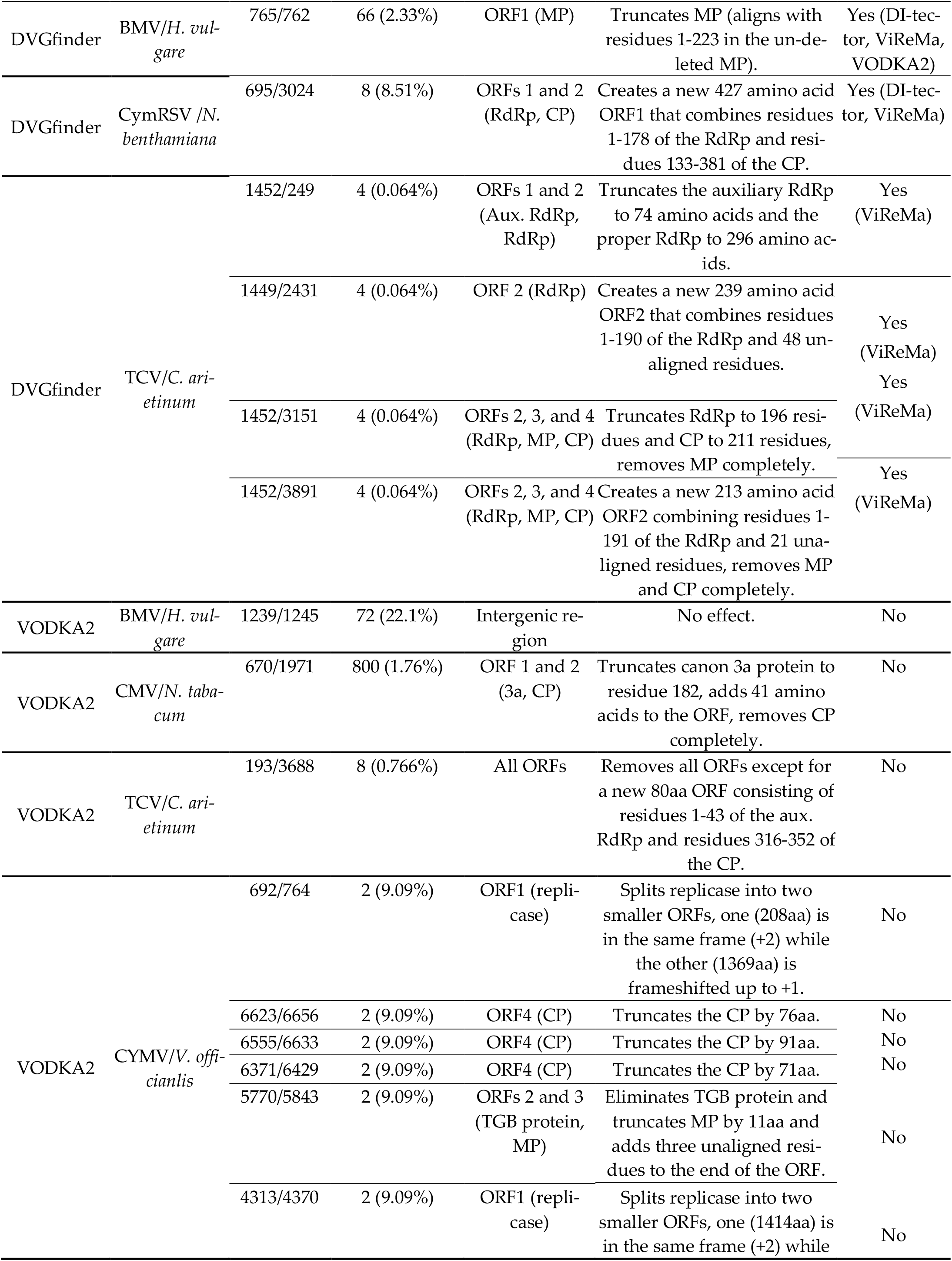
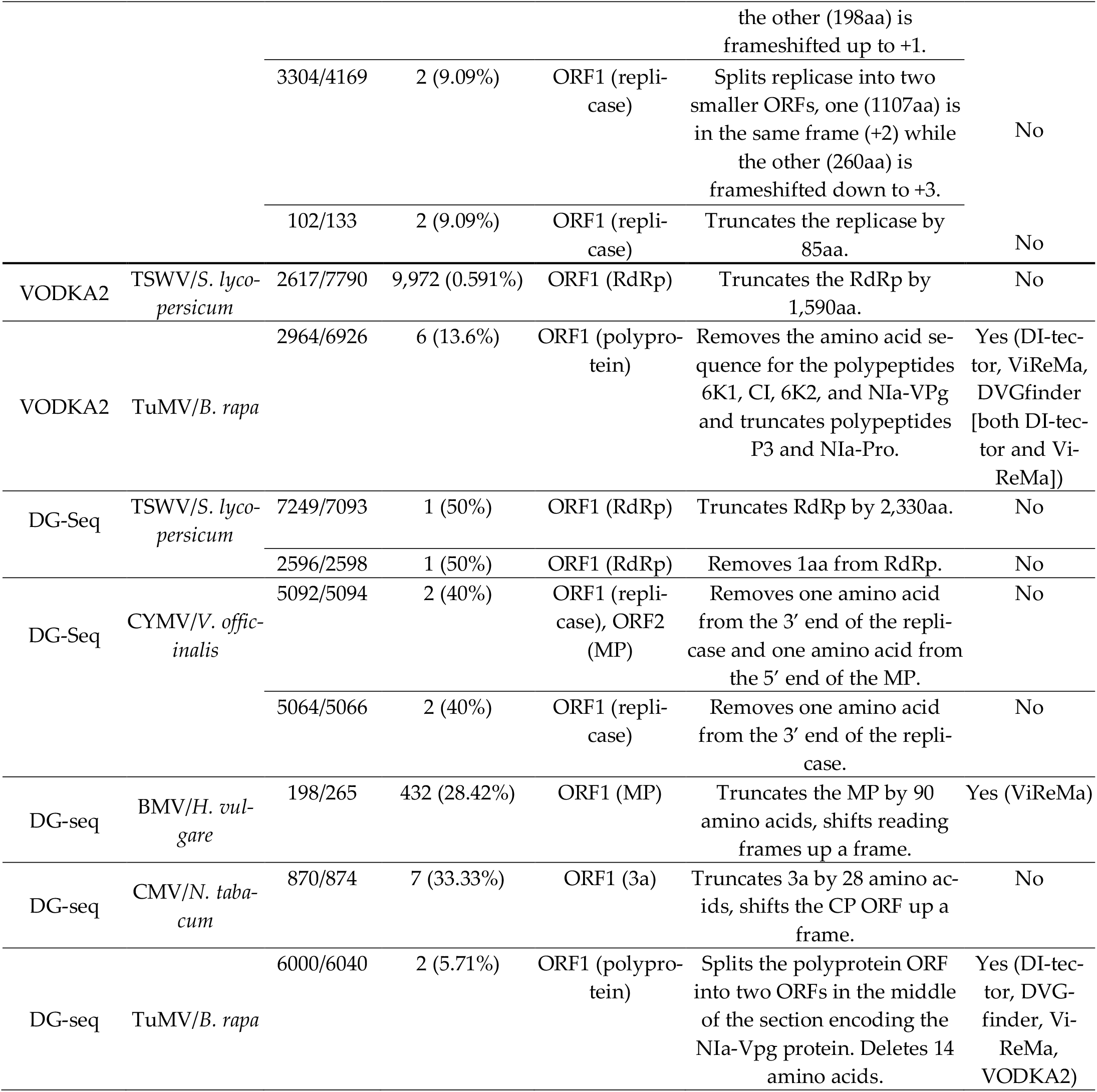
Report on the most common junction site reported by each bioinformatics program, along with information on % total observations (not total reads, because not all programs report total reads) and information regarding what influence that junction has on genomic organization. Instances where no junctions were found more than 1 time were not counted, as were lists of junctions greater than ten with tied hits. Consult supplemental table 2 for a list of what junctions went into a bin and supplemental table 3 for the full list of junctions greater than ten with tied hits. MP = movement protein, CP = coat protein, RdRp = RNA-dependent RNA polymerase, SS = silencing suppressor.

**Figure 1.**
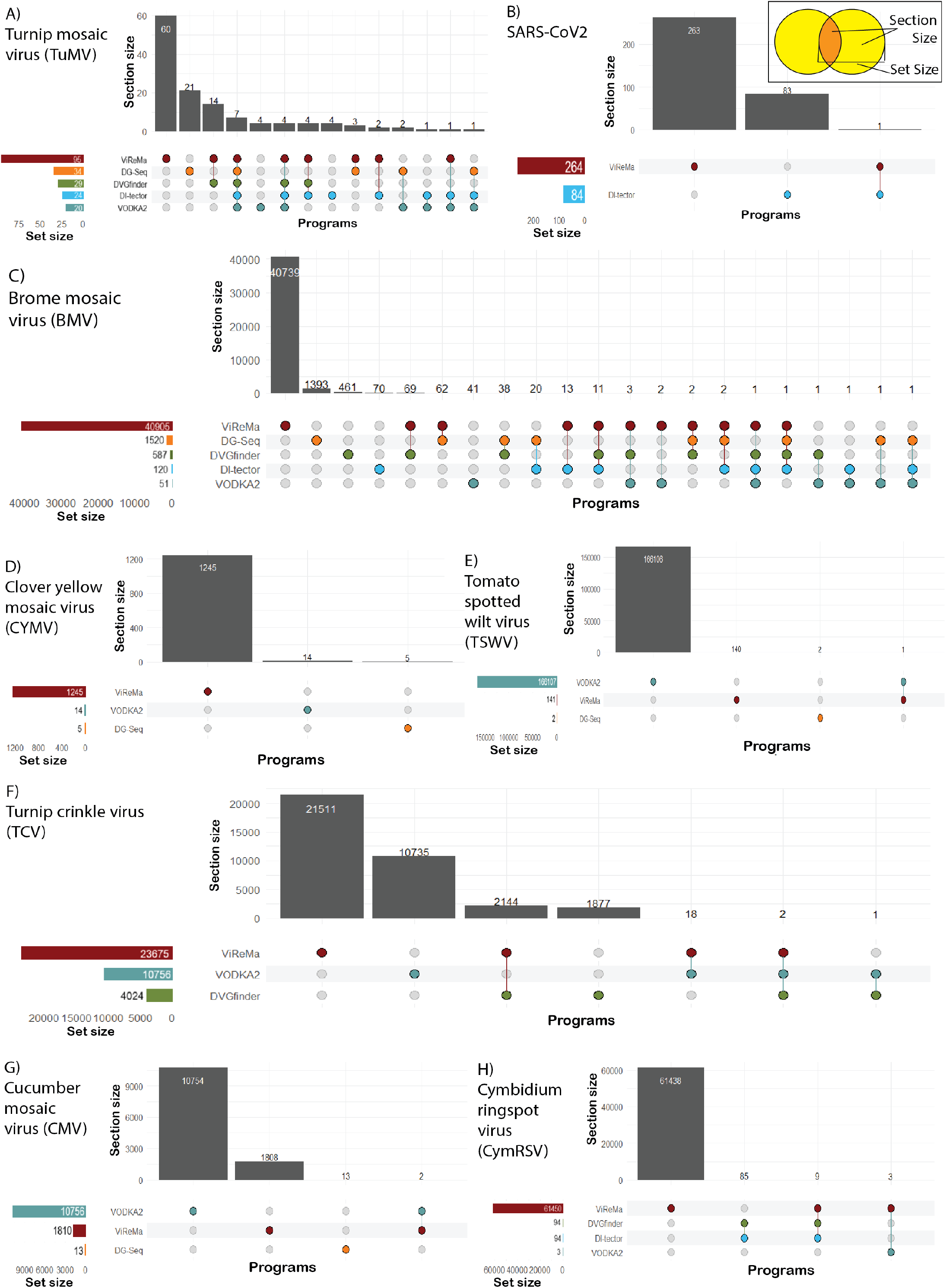
UpSet plots demonstrating the degree of overlap between programs regarding junctions identified and written to their outputs. BMV (C) showed the most complex degree of overlaps, while CYMV (D) showed no overlaps. Control datasets are TuMV (A) and SARS-CoV2 (B).

### 3.2 Different bioinformatics programs do not identify the same junction as the most common, although multiple programs may identify the same junction in their output

The second major question we addressed was whether the outputs from the different bioinformatic tools agreed when the same dataset was run through them. **Fig. 1** soundly demonstrates that for the most part the outputs do not agree, however, we also wanted to know if those that do all identify the same junction as the most common. **Table 3** demonstrates that no, the most commonly occurring junction site(s) identified by different programs also generally does not agree. The only junction site identified as the most common between two programs was 2964/6926 from TuMV, which was identified as the most common in that dataset’s DI-tector output as well as DVGfinder. However, since DVGfinder is not a unique search algorithm (made up instead of both DI-tector and ViReMa with novel downstream processing and unified language), we conclude that this instance is not enough to say that different programs “sometimes” list the same junction site as the most common, since DVGfinder listed this one due to the fact that it was the most common in the DI-tector portion of search algorithm. Therefore, this junction appeared twice as the most common solely due to DI-tector.

However, we did pick up on a handful of junctions that were identified as most common from one program in the outputs of others, they were just not identified as most common in those other outputs. As **table 3** demonstrates, this is mostly due to DVGfinder overlaping with DI-tector and ViReMa, which is to be expected considering DVGfinder’s design. However, two BMV junctions, 786/794 and 765/662, were independently found by DI-tector, ViReMa, and VODKA2, and the BMV junction 198/265 was independently found by ViReMa. This also occurred with the TuMV control junctions 2964/6926 and 6000/6040, and the SARS-CoV2 control junction 60/28250, which was independently found by DI-tector and ViReMa but not DVGfinder.

### 3.4. Some identified DVGs would be worth exploring downstream for interfering capability, while others are most likely neutral mutations

The primary driving force behind using bioinformatics programs to look at the defective portions of the virome is the discovery and characterization of defective interfering RNA (or DNA) that can be used as an antiviral against a virus of interest. Obviously, assessment for interfering capability of a DVG identified in a bioinformatics workflow must be done downstream in laboratory setting, but in many circumstances thousands of unique DVGs are identified in the output (**table 2**). As previously stated, many known defective interfering RNAs are present in high enough abundance that they can be seen as separate bands on a Northern blot [19–24]. Therefore, the most common junction point could correspond to a DI and be an effective starting point for downstream testing. We assessed whether the most common junction point consistently showed promise for downstream validation by taking the most common junction points in **table 3**, re-creating the deletions in the viral reference genome in SnapGene (v. 7.2.1), and observing the effects on the ORFs of that RNA. The most common deletions typically corresponded to ORF truncation or deletion, sometimes substantially so. However, some common junctions, such as 1239/1245 found by VODKA2 in BMV and 2596/2598 found by DG-Seq in TSWV, had no substantial effect *in silico*.

## 4. Discussion

### 4.1. Our results and DI prevalence theory

Decisively answering whether bioinformatics programs will reliably be able to find plant virus DI candidates in RNAseq data is hampered by the fact that much of what leads to the evolution and maintenance of DIs in virus infected plants is a black box. The simple answer to whether bioinformatics programs can identify DIs is no: the only thing that they can do is identify when a read is a chimera of more than one contiguous sequence in the given reference. Without additional context, that information cannot be taken as evidence of a DI, or even a DVG. However, because in many cases we *have* additional context, the answer to this question is slightly more complicated.

As previously stated, our working hypothesis regarding DI prevalence in virus populations is that as long as virally encoded proteins can assist in genome replication in *trans* and that these *trans-*assisting virally encoded proteins increase the fitness of the genome they help replicate non-linearly, DIs should be present in the viral quasispecies. However, there are undoubtably other as-of-yet-unknown variables in the circumstances that influence DI generation and the maintenance and/or accumulation of specific DI genome organizations in different virus/host combinations. Regardless, this hypothesis leads to the prediction that if DIs are found in one instance of a virus/host combination, then the conditions for DI generation and accumulation---whatever they are---are permissive and we should still be able to locate DIs of that particular genome organization in any other instance of the same combination of virus species/host species.

We assessed the validity of this prediction to the best of our ability in our experimental design, by choosing three datasets that came from a virus/host combination that has been previously shown in the literature to generate DIs. The prediction seemed to hold in 2/3 cases, however, the third dataset, CMV/*N. tabacum*, did not provide evidence to support this. This could be explained by the fact that this dataset didn’t perfectly match the parameters of our prediction: the strain of CMV used was different that the strain previously published to produce DIs. However, strain is a below-species classification of viruses, so the fact that the CMV dataset failed to support our prediction may indicate that our prediction is indeed false and ought to be narrowed to the prediction of “DIs of a known genome organization should arise in any other instance of the same combination of virus/host *genotypes*” rather than species.

Given our working hypothesis, we also predicted that we likely wouldn’t find previously published DI junctions in novel hosts, since the fitness landscape of the viral population in that host would lead to DIs of a different genome organization, if that host is permissive. However, one of the virus/novel host datasets, TCV in *C. arientum*, had a canon junction site located in ViReMa’s output, and only one instance of it. Considering that previously research on TCV DIs find them in high enough abundance to get clear Northern blots [21], and ViReMa does produce false positives [26], we consider a single instance of that junction, only identified by one program, a false positive. To play devil’s advocate, this dataset was of siRNA, and previous work has demonstrated that tombusvirus DIs appear to be poor targets for siRNA generation [2]. Since TCV is also a tombusvirus, it is possible that there were plentiful DIs in this infection but that they are not reflected in the siRNA population. A definitive answer as to whether *C. arientum* infected with TCV-B (NC_003821.3) produces DIs will require wet-lab validation.

### 4.2. Variability in program output

Running the same dataset through different programs and getting some differences in output has been previously noted between ViReMa and DG-seq [28]. Ergo, our results to this end are consistent with the literature, although we are the only paper to our knowledge to compare outputs from a single dataset across *all* currently available bioinformatics pipelines. The results of this, as **figure 1** shows, is that there’s little overlap between program outputs: even DVGfinder does not overlap completely with DI-tector and ViReMa outputs when those two are run independently. This lack of consensus does not instill confidence that these programs are useful, but this is not necessarily an analysis drawback, for it re-affirms the need to rigorously characterize a quasispecies through wetlab work, a sentiment that may have been forgotten with the buzz and novelty of high computing power and cheap sequencing. Running one dataset through multiple programs also provides different “points of view,” which can be used to generate research questions to be explored through empirical experiments. It is easy to hypothesize that junctions with a higher degree of program consensus have a higher likelihood of being “real” (e.g., a junction site found in just two programs vs. five is more likely to be an artifact of sequencing or analysis), however, this hypothesis will have to be addressed using sequencing data specifically taken from an experiment to test this.

### 4.3. Analysis drawbacks

The analysis presented here does have several weak points that should be addressed. For one, although comparing datasets from multiple labs does adequately address our hypothesis of DI prevalence, our conclusions would be more robust if all the datasets were the same type of RNAseq. This, coupled with the fact that they are from entirely different research groups, means that they are not directly comparable. Our hypotheses did not require a direct comparison, but all our claims would greatly benefit from sequencing experiments specifically designed to address our questions.

Furthermore, one major drawback of using NGS for this kind of DVG and/or DI investigation is the inability to identify small mosaic genomes, which, at least in plants, tend to be the DIs that alter infection dynamics enough to be noticed and characterized in all the previously cited literature. To identify these genomes, one needs long-read sequencing data, of which our SARS-CoV2 dataset was, but long-read sequencing data does not appear to play nice with bioinformatics programs currently available for DVG/DI discovery, with only a BWA-MEM alignment of the reads to the sequence of the junctions locating the junction points (**table 2**).

Furthermore, on the topic of controls, it is interesting to note that even the *in silico* generated dataset failed to have 100% consensus across all programs. In fact, for the TuMV dataset, DG-seq found the most instances of the purposely seeded DIs, at 34, beating out even the program the dataset was designed for, DVGfinder (**table 2**).

## 5. Conclusions

In this paper, we explored the utility of currently available bioinformatics programs for identifying junction points in HTS data that could be tested downstream for antiviral capacity. Specifically, we looked at whether the outputs from these bioinformatic tools generally agree and whether the most frequently identified DVGs from these programs are promising DI candidates for downstream validation. We also explored the possibility of these tools helping us address a larger research question of whether DI RNA are consistently generated and maintained in a specific virus-host combination when conditions are permissive for their replication and accumulation, our “DI prevalence” hypothesis. Our results demonstrate a low degree of agreement regarding identified junction points between programs, promise regarding looking at the most commonly occurring junction for DI candidates, and support for our DI prevalence hypothesis. Bioinformatics workflows have a place in the toolbox of DI and DVG research, but they should not be used alone. We suggest the use of multiple programs on a dataset to better inform decisions regarding deletions to re-create and screen downstream and reiterate the importance of other avenues of evidence in DVG/DI characterization.

## Author Contributions

Conceptualization, A.A.T.; A.A.T.; validation, A.A.T.; formal analysis, A.A.T.; investigation, A.A.T.; data curation, A.A.T.; visualization, A.A.T.; writing—original draft preparation, A.A.T.; writing—review and editing, A.A.T., M.A., and C.R.; supervision, M.A. and C.R. All authors have read and agreed to the published version of the manuscript.

## Data Availability Statement

All data can be found following the links or SRA/GenBank accession numbers provided in the materials and methods section of this paper. All code used for processing bioinformatic program outputs can be found at https://github.com/skybird99-anthony-taylor/DIs_in_RNAseq_data.

## Acknowledgment

E. Achouri, for collaboration on re-writing VODKA2 code; A. Routh, for assistance troubleshooting ViReMa; M. Olmo-Uceda, for assistance troubleshooting DVGfinder; and Molly-Mae M. Conrado and Amelia J. Taylor, for assistance in data analysis.

## Disclaimer/Publisher’s Note

The statements, opinions and data contained in all publications are solely those of the individual author(s) and contributor(s) and not of MDPI and/or the editor(s). MDPI and/or the editor(s) disclaim responsibility for any injury to people or property resulting from any ideas, methods, instructions or products referred to in the content.

## References

1. Rand, U.; Kupke, S.Y.; Shkarlet, H.; Hein, M.D.; Hirsch, T.; Marichal-Gallardo, P.; Cicin-Sain, L.; Reichl, U.; Bruder, D. Antiviral Activity of Influenza a Virus Defective Interfering Particles against Sars-Cov-2 Replication in Vitro through Stimulation of Innate Immunity. Cells 2021, 10, doi:10.3390/cells10071756.

2. Szittya, G.; Molnár, A.; Silhavy, D.; Hornyik, C.; Burgyán, J. Short Defective Interfering RNAs of Tombusviruses Are Not Targeted but Trigger Post-Transcriptional Gene Silencing against Their Helper Virus. Plant Cell 2002, 14, 359–372, doi:10.1105/tpc.010366.

3. Id, A.L.; Young, P.G.; Turner, P.E.; Wild, G.; West, A. Cheating Leads to the Evolution of Multipartite Viruses. PLoS Biol 2023, 21, 1–24, doi:10.1371/journal.pbio.3002092.

4. von Magnus, P. Incomplete Forms of Influenza Virus. Adv Virus Res 1954, 2, 59–79.

5. Hillman, B.I.; Carrington, J.C.; Morris, J.T. A Defective Interfering RNA That Contains a Mosaic of a Plant Virus Genome. Cell 1987, 51, 427–433, doi:10.1016/0092-8674(87)90638-6.

6. Scholthof, K.B.G.; Scholthof, H.B.; Jackson, A.O. The Effect of Defective Interfering RNAs on the Accumulation of Tomato Bushy Stunt Virus Proteins and Implications for Disease Attenuation. Virology 1995, 211, 324–328, doi:10.1006/viro.1995.1410.

7. Adam, G.; Gaedigk, K.; Mundry, K.W. Alterations of a Plant Rhabdovirus during Successive Mechanical Transfers. Journal of Plant Diseases and Protection 1983, 90, 28–35, doi:https://www.jstor.org/stable/43382909.

8. Knorr, D.A.; Mullin, R.H.; Hearne, P.Q.; Morris, J. De Novo Generation of Defective Interfering RNAs of Tomato Bushy Stunt Virus by High Multiplicity Passage. Virology 1991, 181, 193–202, doi:10.1016/0042-6822(91)90484-S.

9. Celix, A.; Rodriguez-Cerezo, E.; Garcia-Arenal, F. New Satellite RNAs, but No DI RNAs, Are Found in Natural Populations of Tomato Bushy Stunt Tombusvirus. Virology 1997, 239, 277–284, doi:10.1006/viro.1997.8864.

10. Budzynska, D.; Zwart, M.P.; Hasiow-Jaroszewska, B. Defective RNA Particles of Plant Viruses — Origin, Structure and Role in Pathogenesis. Viruses 2022, 14, 2814.

11. Law, M.D.; Morris, J.T. De Novo Generation and Accumulation of Tomato Bushy Stunt Virus Defective Interfering RNAs without Serial Host Passage. Virology 1994, 198, 377–380.

12. Budzyńska, D.; Minicka, J.; Hasiów-Jaroszewska, B.; Elena, S.F. Molecular Evolution of Tomato Black Ring Virus and de Novo Generation of a New Type of Defective RNAs during Long-Term Passaging in Different Hosts. Plant Pathol 2020, 69, 1767–1776, doi:10.1111/ppa.13258.

13. Mawassi, M.; Gafny, R.; Gagliardi, D.; Bar-Joseph, M. Populations of Citrus Tristeza Virus Contain Smaller-than-Full-Length Particles Which Encapsidate Sub-Genomic RNA Molecules. Journal of General Virology 1995, 76–651.

14. Mawassi, M.; Karasev, A. V; Mietkiewska, E.; Gafny, R.; Lee, R.F.; Dawson, W.O.; Bar-Joseph, M. Defective RNA Molecules Associated with Citrus Tristeza Virus. Virology 1995, 208, 383–387.

15. Romero, J.; Huang, Q.; Pogany, J.; Bujarski, J.J. Characterization of Defective Interfering Rna Components That Increase Symptom Severity of Broad Bean Mottle Virus Infections. Virology 1993, 194, 576–584, doi:10.1006/viro.1993.1297.

16. Stenger, D.C. Genotypic Variability and the Occurrence of Less than Genomic-Length Viral DNA Forms in a Field Population of Beet Curly Top Geminivirus. Phytopathology 1995, 85, 1316–1322.

17. Visser, P.B.; Brown, D.J.F.; Brederode, F.T.; Bol, J.F. Nematode Transmission of Tobacco Rattle Virus Serves as a Bottleneck to Clear the Virus Population from Defective Interfering RNAs. Virology 1999, 263, 155–165, doi:10.1006/viro.1999.9901.

18. Albiach-Martí, M.R.; Guerri, J.; Hermoso De Mendoza, A.; Laigret, F.; Ballester-Olmos, J.F.; Moreno, P. Aphid Transmission Alters the Genomic and Defective RNA Populations of Citrus Tristeza Virus Isolates. Phytopathology 2000, 90, 134–138, doi:10.1094/PHYTO.2000.90.2.134.

19. Burgyan, J.; Grieco, F.; Russo, M. A Defective Interfering RNA Molecule in Cymbidium Ringspot Virus Infections. Journal of General Virology 1989, 70, 235–239, doi:10.1099/0022-1317-70-1-235.

20. Inoue-Nagata, A.K.; Kormelink, R.; Nagata, T.; Kitajima, E.W.; Goldbach, R.; Peters, D. Effects of Temperature and Host on the Generation of Tomato Spotted Wilt Virus Defective Interfering RNAs. Virology 1997, 87, 1168– 1173.

21. Li, X.H.; Heaton, L.A.; Morris, T.J.; Simon, A.E. Turnip Crinkle Virus Defective Interfering RNAs Intensify Viral Symptoms and Are Generated de Novo. Proc Natl Acad Sci U S A 1989, 86, 9173–9177, doi:10.1073/pnas.86.23.9173.

22. White, K.A.; Bancroft, J. 6; Mackie, G.A. Defective RNAs of Clover Yellow Mosaic Virus Encode Nonstructural/Coat Protein Fusion Products. Virology 1991, 183, 479–486.

23. de Oliveira Resende, R.; De Haan, P.; Van De Vossen, E.; De Avila, A.C.; Goldbach, R.; Peters, D. Defective Interfering L RNA Segments of Tomato Spotted Wilt Virus Retain Both Virus Genome Termini and Have Extensive Internal Deletions. Journal of General Virology 1992, 73, 2509–2516, doi:10.1099/0022-1317-73-10-2509.

24. de Oliveira Resende, R.; de Haan, P.; de Avila, A.C.; Kitajima, E.W.; Kormelink, R.; Goldbach, R.; Peters, D. Generation of Envelope and Defective Interfering RNA Mutants of Tomato Spotted Wilt Virus by Mechanical Passage. Journal of General Virology 1991, 72, 2375–2383, doi:10.1099/0022-1317-72-10-2375.

25. González Aparicio, L.J.; López, C.B.; Felt, S.A. A Virus Is a Community: Diversity within Negative-Sense RNA Virus Populations. Microbiology and Molecular Biology Reviews 2022, 86, doi:10.1128/mmbr.00086-21.

26. Routh, A.; Johnson, J.E. Discovery of Functional Genomic Motifs in Viruses with ViReMa-a Virus Recombination Mapper-for Analysis of next-Generation Sequencing Data. Nucleic Acids Res 2014, 42, 1–10, doi:10.1093/nar/gkt916.

27. Beauclair, G.; Mura, M.; Combredet, C.; Tangy, F.; Jouvenet, N.; Komarova, A. V. DI-Tector: Defective Interfering Viral Genomes’ Detector for next-Generation Sequencing Data. Rna 2018, 24, 1285–1296, doi:10.1261/rna.066910.118.

28. Boussier, J.; Munier, S.; Achouri, E.; Meyer, B.; Crescenzo-Chaigne, B.; Behillil, S.; Enouf, V.; Vignuzzi, M.; Van Der Werf, S.; Naffakh, N. RNA-Seq Accuracy and Reproducibility for the Mapping and Quantification of Influenza Defective Viral Genomes. RNA 2020, 26, 1905–1918, doi:10.1261/rna.077529.120.

29. Olmo-Uceda, M.J.; Muñoz-Sánchez, J.C.; Lasso-Giraldo, W.; Arnau, V.; Díaz-Villanueva, W.; Elena, S.F. DVGfinder: A Metasearch Tool for Identifying Defective Viral Genomes in RNA-Seq Data. Viruses 2022, 14, 1–18, doi:10.3390/v14051114.

30. Bosma, T.J.; Karagiannis, K.; Santana-Quintero, L.; Ilyushina, N.; Zagorodnyaya, T.; Petrovskaya, S.; Laassri, M.; Donnelly, R.P.; Rubin, S.; Simonyan, V.; et al. Identification and Quantification of Defective Virus Genomes in High Throughput Sequencing Data Using DVG-Profiler, a Novel Post-Sequence Alignment Processing Algorithm. PLoS One 2019, 14, doi:10.1371/journal.pone.0216944.

31. Achouri, E.; Felt, S.A.; Hackbart, M.; Rivera-Espinal, N.S.; Lopez, C.B. VODKA2: A Fast an Accurate Method to Detect Non-Standard Viral Genomes from Large RNA-Seq Datasets. RNA 2023, doi:10.1261/rna.079747.123.

32. Sun, Y.; Kim, E.J.; Felt, S.A.; Taylor, L.J.; Agarwal, D.; Grant, G.R.; Lopez, C.B. A Specific Sequence in the Genome of Respiratory Syncytial Virus Regulates the Generation of Copy-Back Defective Viral Genomes. PLoS Pathog 2019, 15, doi:10.1371/journal.ppat.1007707.

33. Graves, M.V; Roossinck, M.J. Characterization of Defective RNAs Derived from RNA 3 of the Fny Strain of Cucumber Mosaic Cucumovirus. J Virol 1995, 69, 4746–4751, doi:10.1128/jvi.69.8.4746-4751.1995.

34. Dexheimer, S.; Shrestha, N.; Chapagain, B.S.; Bujarski, J.J.; Yin, Y. Characterization of Variant RNAs Encapsidated during Bromovirus Infection by High-Throughput Sequencing. Pathogens 2024, 13, doi:10.3390/pathogens13010096.

35. Damayanti, T.; Nagano, H.; Mise, K.; Furusawa, I.; Okuno, T. Brome Mosaic Virus Defective RNAs Generated during Infection of Barley Plants. Journal of General Virology 1999, 80, 2511–2518.

36. Damayanti, T.A.; Nagano, H.; Mise, K.; Furusawa, I.; Okuno, T. Positional Effect of Deletions on Viability, Especially on Encapsidation, of Brome Mosaic Virus D-RNA in Barley Protoplasts. Virology 2002, 293, 314–319, doi:10.1006/viro.2001.1276.

37. Szittya, G.; Moxon, S.; Pantaleo, V.; Toth, G.; Pilcher, R.L.R.; Moulton, V.; Burgyan, J.; Dalmay, T. Structural and Functional Analysis of Viral SiRNAs. PLoS Pathog 2010, 6, 1–15, doi:10.1371/journal.ppat.1000838.

38. Ghasemazadeh, A.; Malgorzata ter Haar, M.; Shams-bakhsh, M.; Pirovano, W.; Pantaleo, V. Shannon Entropy to Evaluate Substitution Rate Variation Among Viral Nucleotide Positions in Datasets of Viral SiRNAs. In Viral Metagenomics: Methods and Protocols; Pantaleo, V., Chiumenti, M., Eds.; Springer Methods: Bari, 2018; pp. 187– 197.

39. Lv, J.; Deng, M.; Li, Z.; Zhu, H.; Wang, Z.; Yue, Y.; Yang, Z.; Xu, J.; Jiang, S.; Zhao, W.; et al. Integrative Analysis of the Transcriptome and Metabolome Reveals the Response Mechanism to Tomato Spotted Wilt Virus. Hortic Plant J 2023, 9, 958–970, doi:10.1016/j.hpj.2022.12.008.

40. Nagata, T.; Inoue-Nagata, A.K.; Prins, M.; Goldbach, R.; Peters, D. Impeded Thrips Transmission of Defective Tomato Spotted Wilt Virus Isolates. Phytopathology 2000, 90, 454–459.

41. Inoue-Nagata, A.K.; Kormelink, R.; Sgro, J.-Y.; Nagata, T.; Kitajima, E.W.; Goldbach, R.; Peters, D. Molecular Characterization of Tomato Spotted Wilt Virus Defective Interfering RNAs and Detection of Truncated L Proteins. Virology 1998, 248.

