## Supplemental Table 2 for "Defective But Promising: Evaluating Bioinformatic Pipelines for Utility of Defective Interfering RNA Discovery in Plant Viral Infections"

| Virus | “average” junction | Junctions that went into it | Counts of those junctions | Program |
| --- | --- | --- | --- | --- |
| CYMV | 152/12 | 152/11, 152/13 | 65, 47 | VIREMA |
| CYMV | 152/47 | 152/52, 152/40, 152/48 | 61, 54, 46 | VIREMA |
| CMV | 38/16 | 38/15, 38/17 | 5, 5 | VIREMA |
| COVID | 29,871/29,890 | 29,871/29,895, 29,871/29,894, 29,871/29,893, 29,871/29,892, 29,871/29,891, 29,871/29,890, 29,871/29,889, 29,871/29,888, 29,871/29,887,  29,871/29,886  29,871/29,885 | 2, 2, 2, 2, 2, 2, 2, 2, 2, 2, 2 | VIREMA |
| TCV | Had 94 non-binnable junctions |  | 4 for all | VIREMA |
| TuMV | Had 95 non-binnable junctions |  | 1 for all | VIREMA |
| BMV | 786/794 | 790/797, 788/795, 787/794, 787/795, 786/794, 786/793, 786/796, 786/795, 785/793, 785/795, 785/792, 784/794, 784/792, 783/792, 782/790 | 12, 12, 105, 26, 125, 113, 19, 16, 51, 33, 16, 13, 12, 10, 33 | DI-TECTOR |
| COVID | 60/28250 | 58/28249, 58/28250, 59/28250, 60/28250, 61/28250, 58/28251, 60/28251, 61/28251 | 1, 1, 2, 4, 9, 1, 15, 89 | DI-TECTOR |
| CymRSV | 691/3024 | 680/3022, 681/3023, 691/3025, 691/3023, 692/3023, 694/3022, 695/3023, 695/3022, 696/3024, 697/3024 | 1, 11, 4, 1, 1, 2, 13, 3, 9, 2 | DI-TECTOR |
| BMV | 765/762 | 761/759, 762/759, 766/762, 769/767 | 16, 16, 16, 18 | DVGFINDER |
| CymRSV | 695/3024 | 691/3023,  691/3025,  692/3023,  694/3022,  695/3022,  695/3023,  696/3024,  697/3024 | 1 FOR ALL | DVGFINDER |
| TCV | 1452/249 | 1451/249, 1452/249 | 4 for both | DVGFINDER |
| TCV | 1449/2431 | 1447/2429, 1451/2433 | 4 for both | DVGFINDER |
| TCV | 1452/3151 | 1450/3147, 1454/3154 | 4 for both | DVGFINDER |
| TCV | 1452/3891 | 1452/3886, 1451/3895 | 4 for both | DVGFINDER |
| TCV | 193/3688 | 193/3686, 193/3689 | 4 for both | VODKA2 |
| CYMV | 5092/5094 | 5090/5092,  5093/5095 | 1 for both | DG-SEQ |
| CYMV | 5064/5066 | 5063/5065,  5065/5067 | 1 for both | DG-SEQ |
