## Supplemental Table 3 for "Defective But Promising: Evaluating Bioinformatic Pipelines for Utility of Defective Interfering RNA Discovery in Plant Viral Infections"

| Virus | Program | Junction List | Number of hits |
| --- | --- | --- | --- |
| TuMV | VIREMA | 6924/2962  6225/704  9347/8648  1004/7321  4600/5215  9640/5891  8254/3000  7501/8757  1803/5741  1071/1973  3885/7826  8357/5016  4795/4531  6104/4165  6040/6000  6504/742  1502/4275  5083/7573  3611/4844  3598/3792  7470/8365  8665/4479  7242/3859  5147/3516  4653/4559  8595/5065  918/8831  5149/8484  552/5442  2983/7883  176/490  2964/6926  8650/9349  6000/6040  7574/5084  8581/1436  4846/3613  3859/7242  4560/4654  3002/8256  8834/921  490/176  8779/3087  5016/8357  2366/3648  4022/7019  1784/6060  55/8684  745/6507  8076/3193  3792/3598  4062/2928  8224/2747  706/6227  3516/5147  50/6093  1909/8273  1051/2302  5893/9642  8484/5149  6288/7967  970/3497  1460/2998  762/8125  9177/9776  3537/2677  7710/5359  3497/970  8980/1038  1040/8978  8125/762  7968/6287  3460/7774  1707/7655  2677/3537  163/4932  102/4015  3000/1458  4964/8435  3200/9054  6940/9114  2354/4290  2387/1501  9114/6940  8432/4967  1035/2543  8434/7304  3409/4133  4133/3409  6240/3324  3324/6240  6125/9615  8306/7679  4255/7271  7271/4255 | 1 (for all) |
| TCV | VIREMA | 1177/860  3397/3425  1509/2281  2495/3981  3340/1361  796/1347  3952/233  3214/3844  2986/1727  3425/1340  2782/2429  2871/658  3034/1571  1183/3772  1073/3110  1105/2858  2262/726  3522/1726  2584/3756  342/3461  1124/970  2268/2037  374/982  2356/3324  3324/2693  3282/3668  1682/737  259/3622  2581/2373  2869/2051  501/1496  2419/2685  2930/2655  3276/3493  2991/1100  2091/3077  2687/274  3593/1769  1249/593  2174/3619  1216/2091  2753/1458  1453/32  3268/1559  1670/3013  2090/1862  1841/1237  194/3570  3791/3016  3150/2912  1118/2219  1354/3133  3114/505  1760/3982  419/3437  3608/3828  380/1726  221/2579  2021/1195  1015/1251  2239/659  92/457  1896/3898  1805/2110  3844/1351  3821/3754  1304/176  1382/1988  2838/3461  1634/405  3013/2910  2654/3122  3682/2868  3523/554  3281/3602  745/2471  999/3084  2583/3384  1544/1528  563/1125 | 4 (for all) |
| TuMV | DVGFINDER | 4795/4531  5147/3516  6040/6000  6104/4165  6225/704  6504/742  6924/2962  7242/3859  8254/3000  8357/5016  8595/5065  8665/4479  9640/5891  1051/2302  1784/6060  1909/8273  2366/3648  2964/6926  3002/8256  3516/5147  3859/7242  4022/7019  4560/4654  5893/9642  6000/6040  706/6227  745/6507  8650/9349 | 1 (for all) |
