## Supplemental Table 1 for "Defective But Promising: Evaluating Bioinformatic Pipelines for Utility of Defective Interfering RNA Discovery in Plant Viral Infections"

| Junction | Program |
| --- | --- |
| 1051/2302 | DG-SEQ |
| 1784/6060 | DG-SEQ |
| 1909/8273 | DG-SEQ |
| 1973/1071 | DG-SEQ |
| 2962/6924 | DG-SEQ |
| 3000/8254 | DG-SEQ |
| 3516/5147 | DG-SEQ |
| 3792/3598 | DG-SEQ |
| 3859/7242 | DG-SEQ |
| 4022/7019 | DG-SEQ |
| 4060/2926 | DG-SEQ |
| 4165/6104 | DG-SEQ |
| 4275/1502 | DG-SEQ |
| 4479/8665 | DG-SEQ |
| 4531/4795 | DG-SEQ |
| 4559/4653 | DG-SEQ |
| 4844/3611 | DG-SEQ |
| 49/6092 | DG-SEQ |
| 490/176 | DG-SEQ |
| 5065/8595 | DG-SEQ |
| 5215/4600 | DG-SEQ |
| 5741/1803 | DG-SEQ |
| 5891/9640 | DG-SEQ |
| 6000/6040 | DG-SEQ |
| 6000/6040 | DG-SEQ |
| 704/6225 | DG-SEQ |
| 742/6504 | DG-SEQ |
| 7573/5083 | DG-SEQ |
| 7826/3885 | DG-SEQ |
| 7883/2983 | DG-SEQ |
| 8365/7470 | DG-SEQ |
| 8579/1434 | DG-SEQ |
| 8648/9347 | DG-SEQ |
| 8779/3087 | DG-SEQ |
| 8831/918 | DG-SEQ |
| 1784/6060 | DI-TECTOR |
| 2366/3648 | DI-TECTOR |
| 2964/6926 | DI-TECTOR |
| 4019/7017 | DI-TECTOR |
| 4022/7019 | DI-TECTOR |
| 4166/6105 | DI-TECTOR |
| 4533/4797 | DI-TECTOR |
| 5016/8357 | DI-TECTOR |
| 55/8684 | DI-TECTOR |
| 6000/6040 | DI-TECTOR |
| 745/6507 | DI-TECTOR |
| 8650/9349 | DI-TECTOR |
| 1004/7321 | DVGFINDER |
| 102/4015 | DVGFINDER |
| 1040/8978 | DVGFINDER |
| 1051/2302 | DVGFINDER |
| 1071/1973 | DVGFINDER |
| 1460/2998 | DVGFINDER |
| 1502/4275 | DVGFINDER |
| 163/4932 | DVGFINDER |
| 1707/7655 | DVGFINDER |
| 176/490 | DVGFINDER |
| 1784/6060 | DVGFINDER |
| 1803/5741 | DVGFINDER |
| 1909/8273 | DVGFINDER |
| 1973/1071 | DVGFINDER |
| 2354/4290 | DVGFINDER |
| 2366/3648 | DVGFINDER |
| 2387/1501 | DVGFINDER |
| 2677/3537 | DVGFINDER |
| 2964/6926 | DVGFINDER |
| 2983/7883 | DVGFINDER |
| 3000/1458 | DVGFINDER |
| 3002/8256 | DVGFINDER |
| 3200/9054 | DVGFINDER |
| 3324/6240 | DVGFINDER |
| 3409/4133 | DVGFINDER |
| 3497/970 | DVGFINDER |
| 3516/5147 | DVGFINDER |
| 3537/2677 | DVGFINDER |
| 3598/3792 | DVGFINDER |
| 3611/4844 | DVGFINDER |
| 3792/3598 | DVGFINDER |
| 3859/7242 | DVGFINDER |
| 3885/7826 | DVGFINDER |
| 4016/101 | DVGFINDER |
| 4022/7019 | DVGFINDER |
| 4062/2928 | DVGFINDER |
| 4133/3409 | DVGFINDER |
| 4255/7271 | DVGFINDER |
| 4275/1502 | DVGFINDER |
| 4560/4654 | DVGFINDER |
| 4600/5215 | DVGFINDER |
| 4653/4559 | DVGFINDER |
| 4795/4531 | DVGFINDER |
| 4846/3613 | DVGFINDER |
| 490/176 | DVGFINDER |
| 4964/8435 | DVGFINDER |
| 5083/7573 | DVGFINDER |
| 5147/3516 | DVGFINDER |
| 5149/8484 | DVGFINDER |
| 552/5442 | DVGFINDER |
| 5741/1803 | DVGFINDER |
| 5893/9642 | DVGFINDER |
| 6000/6040 | DVGFINDER |
| 6040/6000 | DVGFINDER |
| 6104/4165 | DVGFINDER |
| 6125/9615 | DVGFINDER |
| 6225/704 | DVGFINDER |
| 6240/3324 | DVGFINDER |
| 6288/7967 | DVGFINDER |
| 6504/742 | DVGFINDER |
| 6924/2962 | DVGFINDER |
| 6940/9114 | DVGFINDER |
| 706/6227 | DVGFINDER |
| 7242/3859 | DVGFINDER |
| 7271/4255 | DVGFINDER |
| 745/6507 | DVGFINDER |
| 7470/8365 | DVGFINDER |
| 7501/8757 | DVGFINDER |
| 7574/5084 | DVGFINDER |
| 762/8125 | DVGFINDER |
| 7710/5359 | DVGFINDER |
| 7826/3885 | DVGFINDER |
| 7968/6287 | DVGFINDER |
| 8076/3193 | DVGFINDER |
| 8125/762 | DVGFINDER |
| 8224/2747 | DVGFINDER |
| 8254/3000 | DVGFINDER |
| 8306/7679 | DVGFINDER |
| 8357/5016 | DVGFINDER |
| 8365/7470 | DVGFINDER |
| 8434/7304 | DVGFINDER |
| 8581/1436 | DVGFINDER |
| 8595/5065 | DVGFINDER |
| 8650/9349 | DVGFINDER |
| 8665/4479 | DVGFINDER |
| 8779/3087 | DVGFINDER |
| 8834/921 | DVGFINDER |
| 8980/1038 | DVGFINDER |
| 9114/6940 | DVGFINDER |
| 9177/9776 | DVGFINDER |
| 918/8831 | DVGFINDER |
| 9640/5891 | DVGFINDER |
| 970/3497 | DVGFINDER |
| 1004/7321 | VIREMA |
| 102/4015 | VIREMA |
| 1035/2543 | VIREMA |
| 1040/8978 | VIREMA |
| 1051/2302 | VIREMA |
| 1071/1973 | VIREMA |
| 1460/2998 | VIREMA |
| 1502/4275 | VIREMA |
| 163/4932 | VIREMA |
| 1707/7655 | VIREMA |
| 176/490 | VIREMA |
| 1784/6060 | VIREMA |
| 1803/5741 | VIREMA |
| 1909/8273 | VIREMA |
| 2354/4290 | VIREMA |
| 2366/3648 | VIREMA |
| 2387/1501 | VIREMA |
| 2677/3537 | VIREMA |
| 2964/6926 | VIREMA |
| 2983/7883 | VIREMA |
| 3000/1458 | VIREMA |
| 3002/8256 | VIREMA |
| 3200/9054 | VIREMA |
| 3324/6240 | VIREMA |
| 3409/4133 | VIREMA |
| 3460/7774 | VIREMA |
| 3497/970 | VIREMA |
| 3516/5147 | VIREMA |
| 3537/2677 | VIREMA |
| 3598/3792 | VIREMA |
| 3611/4844 | VIREMA |
| 3792/3598 | VIREMA |
| 3859/7242 | VIREMA |
| 3885/7826 | VIREMA |
| 4022/7019 | VIREMA |
| 4062/2928 | VIREMA |
| 4133/3409 | VIREMA |
| 4255/7271 | VIREMA |
| 4560/4654 | VIREMA |
| 4600/5215 | VIREMA |
| 4653/4559 | VIREMA |
| 4795/4531 | VIREMA |
| 4846/3613 | VIREMA |
| 490/176 | VIREMA |
| 4964/8435 | VIREMA |
| 50/6093 | VIREMA |
| 5016/8357 | VIREMA |
| 5083/7573 | VIREMA |
| 5147/3516 | VIREMA |
| 5149/8484 | VIREMA |
| 55/8684 | VIREMA |
| 552/5442 | VIREMA |
| 5893/9642 | VIREMA |
| 6000/6040 | VIREMA |
| 6040/6000 | VIREMA |
| 6104/4165 | VIREMA |
| 6125/9615 | VIREMA |
| 6225/704 | VIREMA |
| 6240/3324 | VIREMA |
| 6288/7967 | VIREMA |
| 6504/742 | VIREMA |
| 6924/2962 | VIREMA |
| 6940/9114 | VIREMA |
| 706/6227 | VIREMA |
| 7242/3859 | VIREMA |
| 7271/4255 | VIREMA |
| 745/6507 | VIREMA |
| 7470/8365 | VIREMA |
| 7501/8757 | VIREMA |
| 7574/5084 | VIREMA |
| 762/8125 | VIREMA |
| 7710/5359 | VIREMA |
| 7968/6287 | VIREMA |
| 8076/3193 | VIREMA |
| 8125/762 | VIREMA |
| 8224/2747 | VIREMA |
| 8254/3000 | VIREMA |
| 8306/7679 | VIREMA |
| 8357/5016 | VIREMA |
| 8432/4967 | VIREMA |
| 8434/7304 | VIREMA |
| 8484/5149 | VIREMA |
| 8581/1436 | VIREMA |
| 8595/5065 | VIREMA |
| 8650/9349 | VIREMA |
| 8665/4479 | VIREMA |
| 8779/3087 | VIREMA |
| 8834/921 | VIREMA |
| 8980/1038 | VIREMA |
| 9114/6940 | VIREMA |
| 9177/9776 | VIREMA |
| 918/8831 | VIREMA |
| 9347/8648 | VIREMA |
| 9640/5891 | VIREMA |
| 970/3497 | VIREMA |
| 1051_2302 | VODKA2 |
| 1784_6060 | VODKA2 |
| 1909_8273 | VODKA2 |
| 2365_3647 | VODKA2 |
| 2964_6926 | VODKA2 |
| 3002_8256 | VODKA2 |
| 3516_5147 | VODKA2 |
| 3859_7242 | VODKA2 |
| 4022_7019 | VODKA2 |
| 4166_6105 | VODKA2 |
| 4479_8665 | VODKA2 |
| 4531_4795 | VODKA2 |
| 4560_4654 | VODKA2 |
| 5016_8357 | VODKA2 |
| 5065_8595 | VODKA2 |
| 5892_9641 | VODKA2 |
| 6000_6040 | VODKA2 |
| 706_6227 | VODKA2 |
| 743_6505 | VODKA2 |
| 8649_9348 | VODKA2 |
| 8649_9348 | VODKA2 |
